# DNA sequence encodes the position of DNA supercoils

**DOI:** 10.1101/180414

**Authors:** Sung Hyun Kim, Mahipal Ganji, Jaco van der Torre, Elio Abbondanzieri, Cees Dekker

## Abstract

The three-dimensional structure of DNA is increasingly understood to play a decisive role in gene regulation and other vital cellular processes, which has triggered an explosive growth of research on the spatial architecture of the genome. Many studies focus on the role of various DNA-packaging proteins, crowding, and confinement in organizing chromatin, but structural information might also be directly encoded in bare DNA itself. Here, we use a fluorescence-based single-molecule technique to visualize plectonemes, the extended intertwined DNA loops that form upon twisting DNA. Remarkably, we find that the underlying DNA sequence directly encodes the structure of supercoiled DNA by pinning plectonemes at specific positions. We explore a variety of DNA sequences to determine what features influence pinning, and we develop a physical model that predicts the level of plectoneme pinning in excellent agreement with the data. The intrinsic curvature measured over a range of ~70 base pairs is found to be the key property governing the supercoiled structure of DNA. Our model predicts that plectonemes are likely to localize directly upstream of prokaryotic transcription start sites, and this prediction is experimentally verified *in vitro.* Our results reveal a hidden code in DNA that helps to spatially organize the genome.

## Introduction

Control of DNA supercoiling is of vital importance to cells. Torsional strain imposed by DNA-processing enzymes induces supercoiling of DNA, which triggers large structural rearrangements through the formation of plectonemes^1^. Recent biochemical studies suggest that supercoiling plays an important role in the regulation of gene expression in both prokaryotes^2^ and eukaryotes^3,4^. In order to tailor the degree of supercoiling around specific genes, chromatin is organized into independent topological domains with varying degrees of torsional strain^3,5^. Domains that contain highly transcribed genes are generally underwound whereas inactive genes are overwound^6^. Furthermore, transcription of a gene transiently alters the local supercoiling^3,6,7^, while, in turn, torsional strain influences the rate of transcription^8-10^.

For many years the effect of DNA supercoiling on various cellular processes has mainly been understood as a torsional stress that enzymes should overcome or exploit for their function. More recently, supercoiling has been acknowledged as a key component of the spatial architecture of the genome^11-14^. Here bound proteins are typically viewed as the primary determinant of sequence-specific tertiary structures while intrinsic mechanical features of the DNA are often ignored. However, the DNA sequence influences its local mechanical properties such as bending stiffness, curvature, and duplex stability, which in turn alter the energetics of plectoneme formation at specific sequences^15-17^. Unfortunately, the relative importance of these factors that influence the precise tertiary structure of supercoiled DNA have remained unclear^18^. Various indications that the plectonemic structure of DNA can be influenced by the sequence were obtained from biochemical and structural studies^4,19,20^ as well as from work performed *in silico*^21-24^. However, these studies only examined a handful of specific sequences such as phased poly(A)-tracts and a particular high–curvature sequence rich in poly(A)-tracts, making it difficult to determine if curvature, long poly(A)-tracts, or some other DNA feature drives the sequence– structure relationship.

Here, we study how DNA sequence governs the structure of supercoiled DNA by use of a recently developed single-molecule technique termed ISD (Intercalation-induced Supercoiling of DNA)^25^, which uses intercalating dyes to induce supercoiling as well as to observe the resultant tertiary structures in many DNA molecules in parallel. Plectonemes are directly observable as intensity maxima along the DNA, from which their position along DNA can be extracted (see Fig. 1a and Fig. S1). We find a strong relationship between sequence and plectoneme localization. By examining many different sequences, we systematically rule out several possible mechanisms of the observed sequence dependence. Using a model built on basic physics, we show that a specific form of curvature determines the relative plectoneme stability at different sequences. Application of this model to sequenced genomes reveals a clear biological relevance, as we identify a class of plectonemic hot spots that localize upstream of prokaryotic promoters. Subsequently, we confirm that these sequences pin plectonemes in our single molecule assay, testifying to the predictive power of our model. We also discuss several eukaryotic genomes where plectonemes are localized near promoters with a spacing consistent with nucleosome positioning. Taken together, our data show a clear sequence-supercoiling relationship that led us to develop a model that reveals that plectonemes help regulate gene expression and the spatial organization of the genome.

**Fig. 1.**
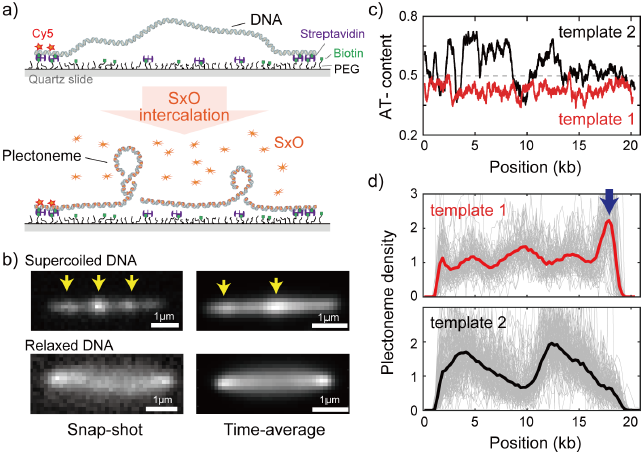
Direct visualization of individual plectonemes on supercoiled DNA. (**a**) Schematic of the ISD assay. (top) A flow-stretched DNA is doubly-tethered on a PEG-coated surface via streptavidin-biotin linkage. One-end of the DNA is labeled with Cy5-fluorophores (red stars) for identifying the direction of each DNA molecule. (bottom) Binding of SxO fluorophores induces supercoiling to the torsionally constrained DNA molecule. (**b**) Representative fluorescence images of a supercoiled DNA molecule. Left: Snap-shot image of a supercoiled DNA with 100ms exposure. Yellow arrows highlight higher DNA density, i.e., individual plectonemes. Right: Time-averaged DNA image by stacking 1000 images (of 100ms exposure each). Arrows indicate peaks in the inhomogeneous average density of plectonemes. (**c**) AT-contents of two DNA samples: template1 and template2 binned to 300-bp. (**d**) Plectoneme densities obtained from individual DNA molecules. (top) Plectoneme density on template1 (grey thin lines, n=70) and their ensemble average (red line). Arrow indicates a strong plectoneme pinning site. (bottom) Plectoneme densities obtained from individual DNA molecules of template2 (grey thin lines, n=120) and their ensemble average (black line).

## Results

### Single-molecule visualization of individual plectonemes along supercoiled DNA

To study the behavior of individual plectonemes on various DNA sequences, we prepared 20 kb-long DNA molecules of which the end regions (~500bp) were labelled with multiple biotins for surface immobilization (Fig. S1b). The DNA molecule were flowed into streptavidin-coated sample chamber at a constant flow rate to obtain stretched double-tethered DNA molecules (Fig. 1a and Fig. S1a). We then induced supercoiling by adding an intercalating dye, Sytox Orange (SxO), into the chamber and imaged individual plectonemes formed on the supercoiled DNA molecules. Notably, SxO does not have any considerable effect on the mechanical properties of DNA under our experimental conditions^25^.

Consistent with previous studies^25,26^, we observed dynamic spots along the supercoiled DNA molecule (highlighted with arrows in Fig. 1b-left and Supp. Movie 1). These spots disappeared when DNA torsionally relaxed upon photo-induced nicking (Fig.1b-bottom)^25^, confirming that the spots were plectonemes induced by the supercoiling. Interestingly, the time-averaged fluorescence intensities of the supercoiled DNA were *not* homogeneously distributed along the molecule (Fig. 1b-top right), establishing that plectoneme occurrence is position dependent. In contrast, torsionally relaxed (nicked) DNA displayed a featureless homogenous time-averaged fluorescence intensity (Fig.1b-bottom right).

### DNA sequence favors plectoneme localization at certain spots along supercoiled DNA

Upon observing the inhomogeneous fluorescence distribution along the supercoiled DNA, we sought to understand if the average plectoneme position is dependent on the underlying DNA sequence. We prepared two DNA samples; the first contained a uniform distribution of AT-bases while the second contained a strongly heterogeneous distribution of AT-bases (Fig. 1c, template1 and template2, respectively). In order to quantitatively analyze the plectoneme distribution, we counted the average number of plectonemes over time at each position on the DNA molecules and built a position-dependent probability density function of the plectoneme occurrence (from now onwards called plectoneme density; see Methods for details). For both DNA samples, we observed a strongly position-dependent plectoneme density (Fig. 1d). Strikingly, the plectoneme densities (Fig. 1d) were very different for the two DNA samples. This difference demonstrates that plectoneme positioning is directed by the underlying DNA sequence. Note that we did not observe any position dependence in the intensity profiles when the DNA was torsionally relaxed, indicating that the interaction of dye is not responsible for the dependence (Fig. S2a).

The plectoneme kinetics showed a similar sequence dependence, as the number of events for nucleation and termination of plectonemes was also found to be position dependent with very different profiles for each DNA samples (Fig. S2b). Importantly, at each position of the DNA, the number of nucleation and termination events were the same, showing that the system was at equilibrium.

### Systematic examination of plectoneme pinning at various putative DNA sequences

We first considered a number of potential links between DNA sequence and plectoneme density. Note that in particular the sharply bent apical tips of plectonemes (Fig. 1A) create an energy barrier to plectoneme formation. This barrier could be reduced if the DNA was able to locally melt or kink, if a specific region of DNA was more flexible than others, or if the DNA sequence was intrinsically curved already before the plectoneme formed. Because all of these properties (duplex stability, flexibility, and curvature) are influenced by the AT-content, we first examined the relationship between AT-content and the measured plectoneme densities in Fig.1c-d. Indeed, the plectoneme density showed a weak correlation with the local AT-percentage (R=0.33, Fig. S3a).

In order to unambiguously link changes in plectoneme density to specific sequences of arbitrary size, we developed an assay where we inserted various short DNA segments carrying particular sequences of interest in the middle of the homogeneous template1 (Fig. 2a and Fig. S3). This allowed us to easily determine the influence of the inserted sequence on plectoneme formation by measuring changes in the plectoneme density at the insert relative to the rest of the DNA strand. We examined three different AT-rich inserts: seqA, seqB, and seqC with ~60%, ~65%, and ~60% AT, respectively (Fig. 2a). Interestingly, all three samples showed a peak in the plectoneme density at the position of insertion, further supporting the idea that AT-rich sequences are preferred positions for plectonemes (Fig. 2b). Furthermore, when we shortened or lengthened one AT-rich sequence (seqA), we found that the probability of plectoneme pinning (i.e. the area under the peak) scaled with the length of the AT-rich fragment (Fig. S3b-e).

**Fig. 2.**
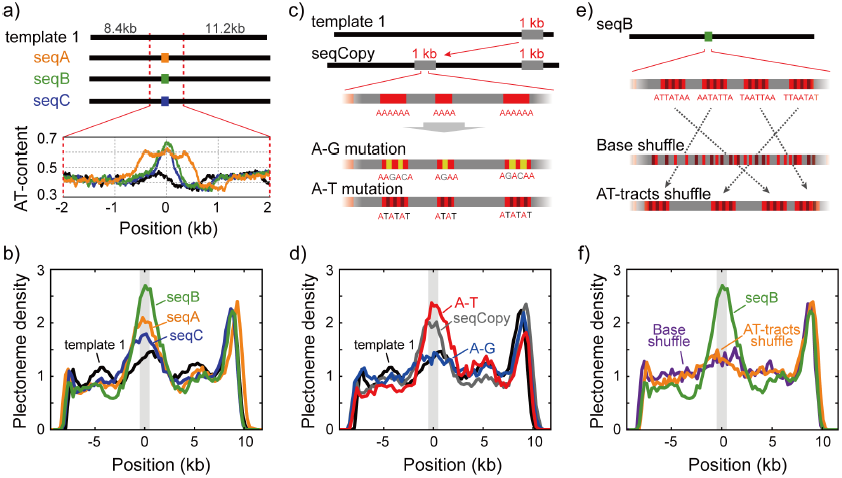
Sequence-dependent pinning of DNA plectonemes. (**a**) Top: Schematics showing DNA constructs with AT-rich fragments inserted in template1. Three different AT-rich segments, SeqA (400bp), SeqB (500bp), and SeqC (1kb), are inserted at 8.8kb from Cy5-end in template1. Bottom: AT-contents of these DNA constructs zoomed in at the position of insertion. (**b**) Averaged plectoneme densities measured for the AT-rich fragments denoted in (A). The insertion region is highlighted with a gray box. (**c**) Schematics of DNA constructs with a copy of the 1kb region near the right end of template1 where strong plectoneme pinning is observed (seqCopy). Poly(A)-tracts within the copied region are then mutated either by replacing A bases with G or C (A-G mutation), or with T (A-T mutation). (**d**) Plectoneme densities measured for the sequences denoted in (c). Plectoneme density of template1 is shown in black, seqCopy in green, A-G mutation in blue, and A-T mutation in red. (**e**) Schematics of DNA constructs with mixed A/T stretches modified from seqB. The insert is modified either by shuffling nucleotides within the insert to destroy all the poly(A) and poly(A/T)-tracts (Base shuffle), or by re-positioning the poly(A) or poly(A/T)-tracts (AT-tracts shuffle) – both while maintaining the exact same AT content across the insert. (**f**) Plectoneme densities measured for the sequences denoted in (e). seqB from panel (b) is plotted in green; base shuffle data are denoted in purple; AT-tracts shuffle in orange.

However, AT-content alone is not the only factor that determines the plectoneme density. For example, the right-end of template1 exhibits a region that pins plectonemes strongly (Fig. 1d-top, arrow), even though this region is not particularly AT-rich (Fig. 1c). When we inserted a 1-kb copy of this pinning region into the middle of template1 (Fig. 2c, ‘seqCopy’), we observed an additional peak in plectoneme density (Fig. 2d, green). Given this region had the same total AT-content as the surrounding DNA, we hypothesized that the distribution of A and T bases may be more important than the total AT-content alone. In particular, poly(A)-tracts influence the local mechanical properties of DNA and might be responsible for the plectoneme pinning, as suggested by early studies^21-23^. To test this, we removed all poly(A) tracts of length 4 or higher by replacing alternative A-bases with G or C-bases in seqCopy (Fig. 2c, ‘A-G mutation’). The peak in the plectoneme density indeed disappeared (Fig. 2d, blue), seemingly confirming our hypothesis. However, when we instead disrupted the poly(A)≥4-tracts by replacing them with alternating AT-stretches (Fig. 2c, ‘A-T mutation’), we, surprisingly, did observe strong pinning (Fig. 2d, red), establishing that plectoneme pinning does not strictly require poly(A)-tracts either. In follow-up experiments, we re-examined the seqB construct to test if long stretches of “weak” bases (i.e. A or T) were the source of pinning. Here, we broke up all poly(A/T)≥4 tracts (i.e. all linear stretches with a random mixture of A or T bases but no G or C bases) by shuffling bases within the seqB insert while keeping the overall AT-content the same. This eliminated plectoneme-pinning, consistent with the idea that poly(A/T) tracts were the cause (Fig. 2e-f, purple). However, if we instead kept all poly(A/T)≥4 tracts intact, but merely rearranged their positions within the seqB insert (again keeping AT-content the same), this rearrangement also abolished the pinning pattern (Fig. 2f, orange).

Taken together, this systematic exploration of various sequences showed that although pinning correlates with AT content, we could not attribute this correlation to AT content alone, to poly(A)-tracts, or to poly(A/T)-tracts. Our data instead suggests that plectoneme pinning depends on the specific distribution of bases, and our shuffled poly(A/T) constructs suggest this distribution must be measured over distances greater than tens of nucleotides. Therefore, among the three mechanical properties we first considered, duplex stability, flexibility, and curvature, the duplex stability is unlikely to be a determinant factor for the plectoneme pinning because duplex stability is mostly determined by the overall AT/GC percentage rather than the specific distribution of bases in the local region.

### Intrinsic local DNA curvature determines the pinning of supercoiled plectonemes

To obtain a more fundamental understanding of the sequence specificity underlying the plectoneme pinning, we developed a novel physical model based on intrinsic curvature and flexibility for estimating the plectoneme energetics (see Methods for details). Briefly, our model estimates the energy cost associated with bending the DNA into the highly curved (~240° arc) plectoneme tip^27^. For example, at 3pN of tension (characteristic for our stretched DNA molecules), the estimated size of the bent tip is 73-bp, and the energy required to bend it by 240° is very sizeable, ~18k_B_T (Fig. 3a). However, if a sequence has a high local intrinsic curvature or flexibility, this energy cost decreases significantly. For example, an intrinsic curvature of 60° between the two ends of a 73-bp segment would lower the bending energy by significant amount, ~8 k_B_T. Hence, we expect that this energy difference drives plectoneme tips to pin at specific sequences. We calculated local intrinsic curvatures at each segment along a relaxed DNA molecule using published dinucleotide parameters for tilt/roll/twist (Fig. 3a and supplement Table 1)^28^. The local flexibility of the DNA was estimated by adding the dinucleotide covariance matrices for tilt and roll^29^ over the length of the loop. Using this approach, we estimate the bending energy of a plectoneme tip centered at each nucleotide along a given sequence (Fig. 3b). The predicted energy landscape is found to be rough with a standard deviation of about ~1 k_B_T, in agreement with a previous experimental estimate based on plectoneme diffusion rates^26^. We then used these bending energies to assign Boltzmann-weighted probabilities, 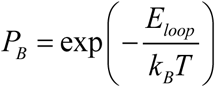, for plectoneme tips centered at each base on a DNA sequence. This provided theoretically estimated plectoneme densities as a function of DNA sequence. Note that we obtained these profiles without any adjustable fitting parameters as the tilt/roll/twist and flexibility values were determined by dinucleotide parameters adopted from published literature. Although both intrinsic curvature and flexibility were included, the model predicts that intrinsic curvature is the most important factor in positioning plectonemes (Fig. 3c).

**Fig. 3.**
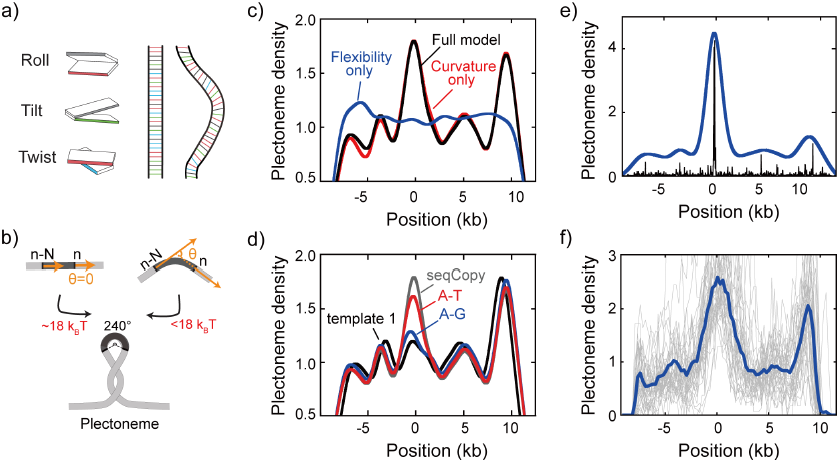
DNA plectonemes pin to sequences that exhibit local curvature. (**a**) Ingredients for an intrinsic-curvature model that is strictly based on dinucleotide stacking. (Left) Cartoons showing the relative alignment between the stacked bases which are characterized by three modes: roll, tilt, and twist. (Middle) In the absence of variations in the roll, tilt, and twist, a DNA molecule adopts a strictly linear conformation in 3D space. (Right) Example of a curved free path of DNA that is determined by the slightly different values for intrinsic roll, tilt, and twist angles for every dinucleotide. (**b**) Schematics showing the energy required to bend a rigid elastic rod as a simple model for the tip of a DNA plectoneme. (**c**) Plectoneme density prediction based on intrinsic curvature and/or flexibility for seqCopy. Predicted plectoneme densities calculated based on either DNA flexibility (blue), only curvature (red), or both (black). Combining flexibility and curvature did not significantly improve the prediction comparing to that solely based on DNA curvature. (**d**) Predicted plectoneme densities for the DNA constructs carrying a copy of the end peak and its mutations, as in Fig. 2b. Note the excellent correspondence to the experimental data in Fig. 2b. (**e-f**) Predicted (e) and measured (f) plectoneme density of a synthetic sequence (250-bp) that is designed to strongly pin a plectoneme. Raw data from the model are shown in black and its Gaussian-smoothed (FWHM=1600bp) is shown in blue in the left panel. Plectoneme densities measured from individual DNA molecules carrying the synthetic sequence (thin grey lines) and their averages (thick blue line) are shown in the right panel.

The predicted plectoneme densities (Fig. 3d) are found to be in excellent agreement with the measured plectoneme densities (Fig. 2d). For example, the non-intuitive mutant sequences tested above (A-G and A-T mutations) are faithfully predicted by the model. More generally, we find that the model qualitatively represented the experimental data for all sequences that were tested. The simplicity of the model and the lack of fitting parameters make this agreement all the more striking.

As a test of the predictive power of our model, we designed a 250 bp-long sequence for which our model *a priori* predicted a high local curvature and strong plectoneme pinning (Fig. 3e). When we subsequently synthesized and measured this construct, we indeed observed a pronounced peak in the plectoneme density (Fig. 3f), demonstrating that the model can be used to identify potential plectoneme pinning sites *in silico*.

### Transcription start sites localize plectonemes in *E. coli*

Given the success of our physical model for predicting plectoneme localization, it is of interest to examine if the model identifies areas of high plectoneme density in genomic DNA that might directly relate to biological functions. Given that our model associates plectoneme pinning with high curvature, we were particularly interested to see what patterns might associate with specific genomic regions. For example, in prokaryotes, curved DNA has been observed to localize upstream of transcription start sites (TSS)^30-32^. In eukaryotes, curvature is associated with the nucleosome positioning sequences found near promoters^33^. However, given that our model requires highly curved DNA over long lengths of ~73 bp to induce plectoneme pinning, it was *a priori* unclear if the local curvature identified at promoter sites is sufficient to strongly influence the plectoneme density.

We first used the model to calculate the plectoneme density profile for the entire *E. coli* genome, revealing plectonemic hot spots spread throughout the genomic DNA (Fig. 4a). Interestingly, we find that a substantial fraction of these hot spots are localized ~100-nucleotides upstream of all the transcription start sites (TSS) associated with confirmed genes in the RegulonDB database (Fig. 4b, red)^34^. We then performed a similar analysis of several other prokaryotic genomes (Fig. 4b). We consistently observe a peak upstream of the TSS, but the size of the peak varied substantially between species, indicating that different organisms rely on sequence-dependent plectoneme positioning to different extents. In one organism (*C. crescentus*), the signal was too weak to detect at all. To experimentally confirm that these sequences represent plectonemic hot spots, we inserted two of these putative plectoneme-pinning sites from *E. coli* into template1. Gratifyingly, we indeed observed a strong pinning effect for these sequences in our single-molecule assay (Fig. 4c-d).

**Fig. 4:**
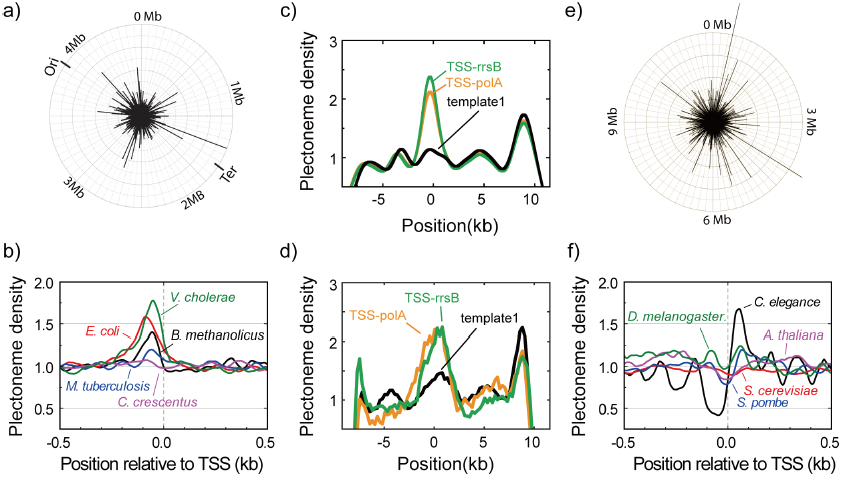
Plectonemes are enriched at prokaryotic transcription start sites. (**a**) The strength of plectoneme pinning calculated for the entire E. coli genome (4,639,221 bp). (**b**) Mean predicted plectoneme densites around transcription start sites (TSS) in prokaryotic genomes. The density profiles were smoothed over a 51 bp window. (**c**) Model-predicted and (**d**) experimentally measured plectoneme densities obtained for two selected TSS sites, TSS-rrsB and TSS-polA, which are E. coli transcription start sites encoding for 16S ribosomal RNA and DNA polymerase I, respectively. For comparison to experimental data, we smoothed the predicted plectoneme densities using a Gaussian filter (FWHM=1600bp) that approximates our spatial resolution. (**e**) Strength of plectoneme pinning calculated for the entire 12.1 Mb genome (i.e. all 16 chromosomes placed in sequential order) of *S. cerevisiae*. For quantitative comparison, we kept the radius of the outer circle the same as in (a). (**f**) Mean predicted plectoneme densities around the most representative TSS for each gene in several eukaryotic genomes (TSS obtained from the Eukaryotic Promoter Database, PMC5210552). The density profiles are smoothed over a 51 bp window.

Finally, we extended our analysis to eukaryotic organisms. Again we found plectonemic hotspots that were spread throughout the genome (Fig. 4e). When averaging near the TSS (Fig. 4f), we found a diverse range of plectoneme positioning signals. While one organism (*S. cervisiae*) showed no detectable plectoneme positioning, most organisms showed both peaks and valleys indicating plectonemes were enriched but also depleted at different regions around the promoter. The features showed a weak periodicity consistent with the reported nucleosome repeat lengths (~150-260 bp)^35^.

## Discussion

In this study, we reported direct experimental observations as well as a novel basic physical model for the sequence-structure relationship of supercoiled DNA. Our single-molecule ISD technique allowed a systematic analysis of sequences that strongly affect plectoneme formation. To explain the underlying mechanism, we developed a physical model that predicts the probability of plectoneme pinning, based solely on the intrinsic curvature and the flexibility of a local region of the DNA. We identified the intrinsic curvature over a ~70bp range as the primary factor that determines plectoneme pinning, while the flexibility alters the mechanics only minimally. Examining full genomes, we found that plectonemes are enriched at promoter sequences in *E. coli* and other prokaryotes, which suggests a role of genetically encoded supercoils in cellular function. Our findings reveal how a previously unrecognized “hidden code” of intrinsic curvature governs the localization of local DNA supercoils, and hence the organization of the three-dimensional structure of the genome.

For a long time, researchers wondered whether DNA sequence may influence the plectonemic structure of supercoiled DNA. Structural and biochemical approaches identified special sequence patterns such as poly(A)-tracts that indicated plectoneme pinning^21-24^. However, the evidence was anecdotal and restricted to a handful of example sequences and it was not possible to establish a general rule for sequence-dependent plectoneme formation. Our high-throughput ISD assay, however, generated ample experimental data that enabled a comprehensive understanding of the underlying mechanism of the sequence-dependent plectoneme pinning.

Our physical modeling reveals that intrinsic curvature is the key structuring factor for determining the three-dimensional structure of supercoiled DNA. In contrast, although perhaps counter-intuitive, we found that the local flexibility is hardly relevant for plectoneme localization. Intrinsic curvatures are encoded in genomic DNA, as evident in our scans of both prokaryotic and eukaryotic genomes, which indicates its biological relevance. In support of this idea, an *in silico* study indeed suggested that curved prokaryotic promoters may control gene expression^36^. Moreover, early *in vivo* studies showed that curved DNA upstream to the promoter site affects gene expression levels^37,38^. These *in vivo* studies suggested that curved DNA would lead to the formation of a small DNA loop, thus facilitating RNAP binding. Our results show that plectonemes are localized on curved DNA, consistent with the presence of a small loop at the tip of the plectoneme right at the curved DNA sequence.

Our analysis of prokaryotic genomes indicates that promoter sequences have evolved local regions with highly curved DNA that promote the localization of DNA plectonemes at these sites. There may be multiple reasons for this. For one, it may help to expose these DNA regions to the outer edge of the dense nucleoid, making them accessible to RNAP, transcription factors, and topoisomerases. Plectonemes may also play a role in the bursting dynamics of gene expression, since each RNAP alters the supercoiling density within a topological domain as it transcribes^6,9^, adding or removing nearby plectonemes^10^. In addition, by bringing distant regions of DNA close together, plectonemes may influence specific promoter-enhancer interactions to regulate gene expression^39^. Finally, plectoneme tips may help RNA polymerase to initiate transcription, since the formation of an open complex also requires bending of the DNA^40^, a mechanism that was proposed as a universal method of regulating gene expression across all organisms^41^. The ability of our model to predict how mutations in the promoter sequence alter the plectoneme density opens up a new way to test these hypotheses.

Our analysis of eukaryotic genomes showed a greater diversity of behavior. The spacing of the peaks suggests that plectonemes may play a role in positioning nucleosomes, consistent with proposals that nucleosome positioning may rely on sequence-dependent signals near promoters^42^. It is also broadly consistent with the universal topological model of plectoneme-RNAP interaction at promoters^41^, which proposes that the plectoneme tip forming upstream of the TSS in eukaryotes is positioned by nearby nucleosomes. The plectoneme signal encoded by intrinsic curvature could therefore indirectly position the promoter plectoneme tip by helping to organize these nearby nucleosomes.

The above findings demonstrate that DNA contains a previously hidden ‘code’ that determines the local intrinsic curvature and consequently governs the locations of plectonemes. These plectonemes can organize DNA within topological domains, providing fine-scale control of the three-dimensional structure of the genome^2^. The model and assay described here make it possible both to predict how changes to the DNA sequence will alter the distribution of plectonemes and to investigate the DNA supercoiling behavior at specific sequences empirically. Using these tools, it will be interesting to explore how changes in this plectoneme code affect levels of gene expression and other vital cellular processes.

## Author contributions

SK, MG, EA, CD conceived the research, SK, MG, JT performed the experiments, SK, MG, EA, and CD analyzed the data, SK, MG, EA, and CD wrote the manuscript.

## Acknowledgments

The data reported in the paper is available from the corresponding authors upon request. We acknowledge valuable discussions with Helmut Schiessel and Ard Louis. We thank Jacob Kerssemakers for helpful discussion and data analysis codes. This work was supported by the ERC Advanced Grant SynDiv [grant number 669598 to C.D.]; the Netherlands Organization for Scientific Research (NWO/OCW) [as part of the Frontiers of Nanoscience program], and the ERC Marie Curie Career Integration Grant [grant number304284 to E.A.].

## Online Methods

### Preparation of DNA molecules of different sequences

Full sequences of all DNA molecules are given in Supplementary Table 2. All DNA molecules except ‘template 2’ in Fig. 1 were prepared by ligating four or five DNA fragments, respectively: 1) ‘Cy5-biotin handle’, 2) ‘8.4-kb fragment’, [ 3) ‘Sequence of Interest’,] 4) ‘11.2-kb fragment’, and 5) ‘biotin handle’ (Fig. S1b). The ‘Cy5-biotin handle’ and ‘biotin handle’ were prepared by PCR methods in the presence of Cy5-modified and/or biotinylated dUTP (aminoallyl-dUTP-Cy5 and biotin-16-dUTP, Jena Bioscience). The ‘8kb-fragment’ and ‘11kb fragment’ were prepared by PCR on Unmethylated Lambda DNA (Promega). These fragments were cloned into pCR-XL using the TOPO^®^ XL PCR cloning kit (Invitrogen) generating pCR-XL-11.2 and pCR-XL-8.4^25^. The fragments were PCR amplified and then digested with BsaI restriction enzyme, respectively. The ‘Sequence of Interest’ was made by PCR on different templates (listed in Supp Table 3). Template 2 in Fig. 1C-black and 1e was made from a digested fragment of an engineered plasmid pSuperCos-λ 1,2 with XhoI and NotI-HF^26^. The digested fragment was further ligated with biotinylated PCR fragments on XhoI side and a biotinylated-Cy5 PCR fragment on the NotI-HF. All the DNA samples were gel-purified before use.

### Dual-color epifluorescence microscopy

Details of our experimental setup are described previously^25,43^. Briefly, a custom-made epifluorescence microscopy equipped with two lasers (532 nm, Samba, Cobolt and 640 nm, MLD, Cobolt) and an EMCCD camera (Ixon 897, Andor) is used to image fluorescently labeled DNA molecules. For the wide-field, epifluorescence-mode illumination on the sample surface, the two laser beams were collimated and focused at the back-focal plane of an objective lens (60x UPLSAPO, NA 1.2, water immersion, Olympus). Back scattered laser light was filtered by using a dichroic mirror (Di01-R405/488/543/635, Semrock) and the fluorescence signal was spectrally separated by a dichroic mirror (FF635-Di02, Semrock) for the SxO channel and Cy5 channel. Two band-pass filters (FF01-731/137, Semrock, for SxO) and FF01-571/72, Semrock, for Cy5) were employed at each fluorescence channel for further spectral filtering. Finally, the fluorescence was imaged on the CCD camera by using a tube lens (f=200 mm). All the measurements were performed at room temperature.

### Intercalation-induced supercoiling of DNA (ISD)

A quartz slide and a coverslip were coated with polyethlyleneglycol (PEG) to suppress nonspecific binding of DNA and SxO. 2% of the PEG molecules were biotinylated for the DNA immobilization. The quartz slide and coverslip were sandwiched with a double-sided tape such that a 100 μm gap between the slide and coverslip forms a shallow sample chamber with flow control. Two holes serving as the inlet and outlet of the flow were placed on the slide glass. Typically, a sample chamber holds 10 μl of solution.

Before DNA immobilization, we incubated the biotinylated PEG surface with 0.1 mg/ml streptavidin for 1 min. After washing unbound streptavidin by flowing 100 μl of buffer A (40 mM TrisHCl pH 8.0, 20mM NaCl, and 0.2 mM EDTA), we flowed the end-biotinylated DNA diluted in buffer A into the sample chamber at a flow rate of 50 μl/min. The concentration of the DNA (typically ~10 pM) was empirically chosen to have an optimal surface density for single DNA observation. Immediately after the flow, we further flowed 200 μl of buffer A at the same flow rate, resulting in stretched, doubly tethered DNA molecules (Fig.1a and Fig.S1a) of which end-to-end extension can be adjusted by the flow rate. We obtained the DNA lengths of around 60-70 % of its contour length (Fig.S1c), which corresponds to a force range of 2-4 pN^25^. We noted that SxO does not exhibit any sequence preference when binding to relaxed DNA, allowing us to back out the amount of DNA localized within a diffraction-limited spot from the total fluorescence intensity (Fig. S2b).

After immobilization of DNA, we flowed in 30 nM SxO (S11368, Thermo Fisher) in an imaging buffer consisting of 40 mM Tris-HCl, pH 8.0, 20 mM NaCl, 0.4 mM EDTA, 2 mM trolox, 40 μg/ml glucose oxidase, 17 μg/ml catalase, and 5 % (w/v) D-dextrose. Fluorescence images were taken at 100 ms exposure time for each frame. The 640nm laser was used for illuminated for the first 10 frames (for Cy5 localization), followed by continuous 532nm laser illumination afterwards. From our previous study we noted that SxO locally unwinds DNA and extends the contour length (Fig. S2b), but does not otherwise affect the mechanical properties of the DNA^25^. Based on the same previous work and assuming that each intercalating dye reduces the twist at the local dinucleotide to zero, we estimate that roughly 1 SxO is bound on every 26 base-pairs of DNA. We note that the numbers of plectoneme nucleation and termination events along supercoiled DNA were equal (Fig. S2b), which is characteristic of a system at equilibrium. Furthermore, we verified that increasing the NaCl concentration from 20 mM to 150 mM NaCl did not result in any significant difference in the observed plectoneme density results, indicating that the plectoneme density is not dependent on the ionic strength (Fig. S3f).

### Data analysis

Analysis of the data was carried out using custom-written Matlab routines, as explained in our previous report^25^. Briefly, we averaged the first ten fluorescence images to determine the end positions of individual DNA molecules. We identify the direction of the DNA molecules by 640 nm illumination at the same field of view, which identifies the Cy5-labelled DNA end. Then, the fluorescence intensity of the DNA at each position along the length was summed up from 11 neighboring pixels perpendicular to the DNA at that position. The median value of the pixels surrounding the molecule was used to correct the background of the image. The resultant DNA intensity was normalized to compensate for photo-bleaching of SxO. We recorded more than 300 frames, each taken with a 100 msec exposure time, and built an intensity kymograph by aligning the normalized intensity profiles in time. Supercoiled DNA intensity profiles, i.e. single lines in the intensity kymograph, were converted to DNA-density profiles by comparing the intensity profile of supercoiled DNA to that of the corresponding relaxed DNA. The latter was obtained after the plectoneme measurements by increasing the excitation laser power that yielded a photo-induced nick of the DNA.

The position of a plectoneme is identified by applying a threshold algorithm to the DNA density profile. A median of the entire DNA density kymograph was used as the background DNA density. The threshold was set at 25 % above the background DNA density. Peaks that sustain at least three consecutive time frames (i.e., ≥300 ms) were selected as plectonemes. For each peak, the cumulative intensities of all the pixels in the right and the left-hand sides of the peak along the DNA were calculated. The intensity ratio between the right and left-hand side was used to find the genomic position of the peak (i.e. base pair position) by comparing this ratio with that obtained from the intensities measured after photo-induced nicking of the molecule of which the pixel position is the same with the genomic position under the given constant tension^25^. After identifying all the plectonemes, the probability of finding a plectoneme (plectoneme density) at each position along the DNA in base-pair space was calculated by counting the total number of plectonemes at each position and normalizing with the observation time. The size of each plectoneme was obtained by integrating intensities of the peak and its two neighboring pixels in both directions (5 pixels total). More than 25 DNA molecules were measured for each DNA sample and the averaged plectoneme densities were calculated with a weight given by the observation time of each molecule. The analysis code written in Matlab (The MathWorks, Inc.) is freely available upon request.

### Plectoneme tip-loop size estimation and bending energetics

An important component of our model is to determine the energy involved in bending the DNA at the plectoneme tip. We first estimate the mean size of a plectoneme tip-loop from the energy stored in an elastic polymer with the same bulk features of DNA. For the simplest case, we first consider a circular loop (360°) formed in DNA under tension. The work associated with shortening the end-to-end length of DNA to accommodate the loop is

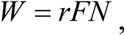

where *F* is the tension across the polymer, *r* is the base pair rise (0.334 nm for dsDNA), and *N* is the number of base pairs. The bending energy is

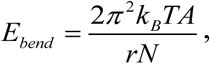

where *k*_*B*_ is the Boltzmann constant, *A* is the bulk persistence length (50 nm for dsDNA). Hence, we obtain an expression for the total energy:

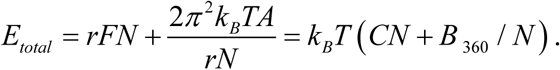

Taking the derivative of *E*_*total*_ with respect to *N* and setting it to zero gives the formula:

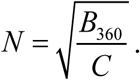

Here, the values of the constants are:

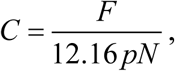

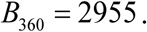

So, at 3 pN we get:

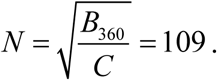

If the loop at the end of the plectoneme is held at the same length but only bent to form a partial circle, the work needed to accommodate the loop will remain the same but the bending energy will be lower, scaling quadratically with the overall bend angle. For a plectoneme tip, a 240° loop is sufficient to match the angle of the DNA in the stem of the plectoneme. The preferred length of a 240° loop is therefore:

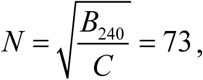

where:

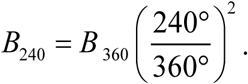

### Physical model predicting the plectoneme density

A full model must explicitly account for the fact that DNA is not a homogeneous polymer. Instead, each DNA sequence has (1) intrinsic curvature and (2) a variable flexibility. Both 1 and 2 depend on the dinucleotide sequences at each location. Note also that we can bend the DNA along any vector normal to the path of the DNA, which describes a circle spanning the full 360° surrounding the DNA strand. We must therefore specify the direction of bending *ϕ* when calculating the bend energy, and we define *ϕ = ϕB* to be the bend direction that aligns with the intrinsic curvature.

The intrinsic curvature can be estimated from the dinucleotide content of the DNA (Fig. 3a). Several studies have attempted to measure the optimal set of dinucleotide parameters (i.e. tilt, roll, and twist) that most closely predict actual DNA conformations^28,44-46^. We find that the parameter set by Balasubramanian et al., produces the closest match to our experimental data when plugged into our model^28^. Using these parameters (see Supp. Table 4), we first calculate the ground state path traced out by the entire DNA strand. We then determine the intrinsic curvature, *θ(N,i)*, across a given stretch of N nucleotides centered at position *i* on the DNA by comparing tangent vectors at the start and end of that stretch. Tangent vectors are calculated over an 11-bp window (1 helical turn, ~3.7 nm). Note that the intrinsic curvature, defined by *θ(N,i)*, also determines the preferred bend direction *ϕB.*

The flexibility of the DNA also varies with position. The flexibility of the tilt and roll angles between neighboring dinucleotide has been estimated by MD simulations^29^. Using these numbers, we can add the roll-tilt covariance matrices for a series of nucleotides (each rotated by the twist angle) to calculate the local flexibility of a given stretch of DNA. The flexibility also depends on the direction of bending. The summed covariance matrix allows us to estimate a local persistence length *A(N,I,ϕ)*.

By combining the local bend angle *θ(N,i)* and the local persistence length *A(N,I,ϕ)*, we are now able to calculate the energy needed to bend a given stretch of DNA to 240°. When the DNA is bent in the preferred curvature direction, this bending energy becomes:

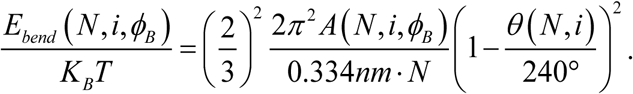

More generally, we can bend the DNA in any direction, in which case the bending energy can be calculated using the law of cosines:

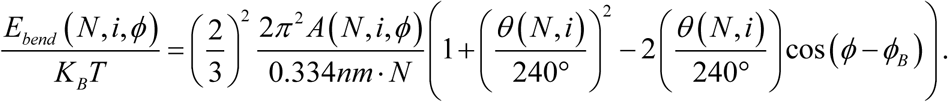

The first formula is the special case when *ϕ = ϕ*_*B*_.

Because both *A(N,i, ϕ)* and *θ(N,i)* are sequence dependent, the loop size and bend direction that minimizes the free energy will also be sequence dependent. Rather than trying to find the parameters that give a maximum likelihood at each position along the template, we find that it is more efficient to calculate the relative probabilities of loops spanning a range of sizes and bend directions. We first calculate the energy associated with each loop using:

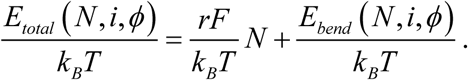

We then assign each of these bending conformations a Boltzmann weight:

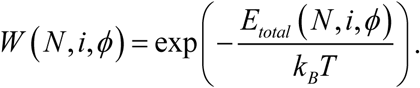

Finally, we sum over all the different bending conformations to get the total weight assigned to the formation of a plectoneme at a specific location *i* on the template:

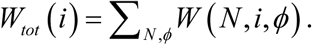

Because the direction *ϕ* is a continuous variable and the length of the loop can range strongly, there are a very large number of bending conformations to sum over. However, because of the exponential dependence on energy, only conformations near the maximum likelihood value in phase space will contribute significantly to the sum. For an isotropic DNA molecule, the maximum likelihood should occur at *N*=73 and *ϕ = ϕ*_*B*_. We therefore sum over parameter values that span this point in phase space. Our final model sums over 8 bending directions (i.e. at every 45°, starting from *ϕ = ϕ*_*B*_.) and calculates loop sizes over a range from 40-bp to 120-bp at 8-bp increments. We verified that the predictions of the model were stable if we increased the range of the loop sizes considered or increased the density of points sampled in phase space, implying that the range of values used was sufficient.

For a fair comparison to experimental data, all predicted plectoneme densities that are presented were smoothened using a Gaussian filter (FWHM=1600bp) that approximates our spatial resolution.

